# Updates to data versions and analytic methods influence the reproducibility of results from epigenome-wide association studies

**DOI:** 10.1101/2021.04.23.441014

**Authors:** Alexandre A. Lussier, Yiwen Zhu, Brooke J. Smith, Andrew J. Simpkin, Andrew D.A.C. Smith, Matthew J. Suderman, Esther Walton, Kerry J. Ressler, Erin C. Dunn

## Abstract

**Introduction:** Biomedical research has grown increasingly cooperative, with several large consortia compiling and sharing epigenomic data. Since data are typically preprocessed by consortia prior to distribution, the implementation of new pipelines can lead to different versions of the same dataset. Analytic frameworks also constantly evolve to incorporate cutting-edge methods and shifting best practices. However, it remains unknown how differences in data and analytic versions alter the results of epigenome-wide analyses, which has broad implications for the replicability of epigenetic associations. Thus, we assessed the impact of these changes using a subsample of the Avon Longitudinal Study of Parents and Children (ALSPAC) cohort.

**Methods:** We analyzed two versions of DNA methylation data, processed using separate preprocessing and analytic pipelines, to examine associations between childhood adversity and prenatal smoking exposure on DNA methylation at age 7. We performed two sets of analyses: (1) epigenome-wide association studies (EWAS); (2) Structured Life Course Modeling Approach (SLCMA), a two-stage method that models time-dependent effects. We also compared results from the SLCMA using more recent methodological recommendations.

**Results:** Differences between ALSPAC data versions impacted both EWAS and SLCMA analyses, yielding different sets of associations at conventional p-value thresholds. However, the magnitude and direction of associations was generally consistent between data versions, regardless of significance thresholds. Updating the SLCMA analytic version similarly altered top associations, but time-dependent effects remained concordant.

**Conclusions:** Changes to data and analytic versions influenced the results of epigenome-wide studies, particularly when using p-value thresholds as reference points for successful replication and stability.

## INTRODUCTION

Biomedical science has become increasingly cooperative over the past decade. The emergence of large datasets, combined with the small effects of biological measures on complex traits, has fueled such cooperation, making global collaboration with researchers more important now than ever. Access to large-scale data has emphasized the importance of identifying both replicable and stable findings, both across and within research studies. As such, large consortia, including birth cohorts, have become an integral part of these collaborative efforts, generating and compiling large amounts of research data ranging from behavioral and clinical markers to molecular and genetic measures. These data are often made available to collaborators and other researchers worldwide, facilitating the interrogation of broader research questions and enabling replication efforts.

Epigenetic data are one key data type collected within these consortia. Epigenetics refer to mechanisms that can result in heritable changes to gene expression without altering genetic sequences ^1^. DNA methylation (DNAm) is the most common type of epigenetic mechanism measured in human studies. DNAm occurs when a methyl residue is added to cytosine residues, typically in the context of cytosine-guanine dinucleotides (CpG). DNAm is both stable over time and responsive to external signals in certain genomic contexts, which highlights its potential as a biomarker and mechanism for the biological embedding of environmental factors ^2^. As such, epigenome-wide association studies (EWAS) have exploded in popularity, with over 1,600 papers on EWAS published since 2015.

To facilitate the sharing of DNAm data, datasets are often processed by the individual cohorts prior to distribution. However, due to both technological and conceptual developments over time, the data available from large cohorts will sometimes become outdated, requiring the distribution of revised versions to collaborators. In addition, individuals in longitudinal studies occasionally withdraw consent to share their data, reducing the overlap of samples between different data versions. At the same time, analytic frameworks are constantly updated and improved upon, resulting in newer cutting-edge methods and shifting analytic best practices ^3^. Yet, the extent to which differences in data versions and analytic pipelines lead to meaningful differences in analytic results remains unclear. This raises an important question as to the replicability and stability of findings across and within studies, which may influence our interpretation of epigenome-wide associations in biomedical research.

Here, we explored the impact of changes in data versions and analytic methods on the consistency of epigenome-wide findings (**Fig 1**). We analyzed two versions of epigenetic data collected from children at age 7 from the Avon Longitudinal Study of Parents and Children (ALSPAC) cohort, a longitudinal birth cohort near Bristol, England. We first characterized the difference between these versions with respect to the distributions of DNAm at the CpG- and individual-level to illuminate the discrepancies that can arise between data versions. Second, we performed two analyses to ascertain the impact of data version changes at the level of CpG-associations, using classical EWAS and a more nuanced analytic method called the Structured Life Course Modeling Approach (SLCMA) ^4^. We performed these analyses using two different types of exposures, contrasting the results from psychosocial (childhood adversity) and physical (maternal smoking during pregnancy) exposures ^5,6^. Finally, we compared results derived from SLCMA between two analytic versions, as more recent guidelines have emerged on its use in big data settings ^3^. Overall, these analyses provide insight into the reproducibility of epigenome-wide associations and highlight the features of epigenetic data that are more reproducible and robust.

**Figure 1.**
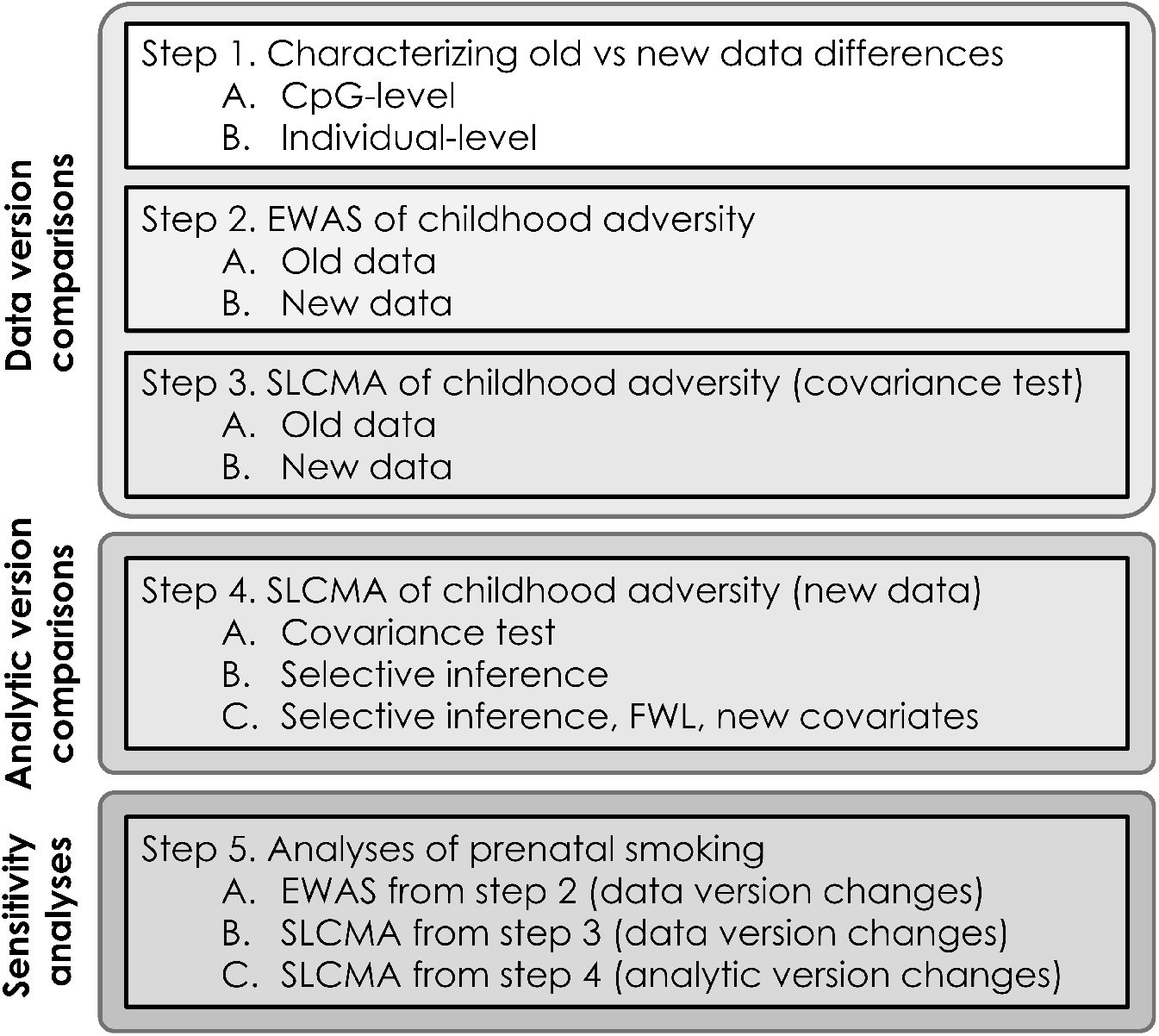
Overview of analyses. Steps 1-3 outline the impact of data version differences. Step 4 outlines the effect of analytic version differences. Here, childhood adversity refers to the seven different types of adversity that were assessed in these analyses. Step 5 outlines the sensitivity analyses of exposure to maternal smoking during gestation, which performed like steps 2-4. *FWL = Frisch-Waugh-Lovell theorem (covariate adjustment methods).

## MATERIALS AND METHODS

### ALSPAC cohort

ALSPAC is a large prospective cohort study that recruited 14,541 pregnancies in Avon, UK, with expected dates of delivery between 1 April 1991 and 31 December 1992 ^7,8^. Further details of the study and available data are provided on the study website through a fully searchable data dictionary (http://www.bris.ac.uk/alspac/researchers/data-access/data-dictionary/). Please note that the study website contains details of all the data that is available through a fully searchable data dictionary and variable search tool (http://www.bristol.ac.uk/alspac/researchers/our-data/). Ethical approval for the study was obtained from the ALSPAC Law and Ethics Committee and the Local Research Ethics Committees. Consent for biological samples has been collected in accordance with the Human Tissue Act (2004). Informed consent for the use of data collected via questionnaires and clinics was obtained from participants following the recommendations of the ALSPAC Ethics and Law Committee at the time. All data are available by request from the ALSPAC Executive Committee for researchers who meet the criteria for access to confidential data (http://www.bristol.ac.uk/alspac/researchers/access/).

### Epigenetic data generation

DNAm profiles at birth, 7, and 15 years of age are part of the Accessible Resource for Integrated Epigenomic Studies (ARIES), a subsample of 1018 mother–child pairs from the ALSPAC cohort ^9^. In this study, we focus on the samples collected at age 7. Briefly, DNA was extracted from peripheral blood samples according to established procedures. DNAm was then measured at 485,577 CpG sites across the genome using the Illumina Infinium Human Methylation 450K BeadChip microarray (Illumina, San Diego, CA). We received two versions of the DNAm data, which were processed using different pipelines by ALSPAC, as described below.

### Epigenetic data versions

In the first version, which we refer to as the *old data* (2015 version), DNAm data were processed using the pipeline developed by Touleimat and Tost ^9,10^. This pipeline involved performing background correction and quantile normalization using the R-package *wateRmelon*. DNAm values for all 485,577 CpGs were provided in the old data version.

In the second data version, which we refer to as the *new data* (2018 version), DNAm data were processed using the pipeline developed by Min and colleagues ^11^. In this version, background correction and functional normalization of DNAm data were performed using the R-package *meffil*. In addition, samples with > 10% of CpG sites with a detection p-value > 0.01 or a bead count < 3 in > 10% of probes were removed. As such, there were fewer CpGs available for analysis (482,855) in the new data compared to the old data (**Fig 2A**). Furthermore, due to data processing and potential removal of consent for some individuals, only 948 participants overlapped between both data versions (**Fig 2A**). Only singleton birth participants present in both data versions were analyzed (n=946).

**Figure 2.**
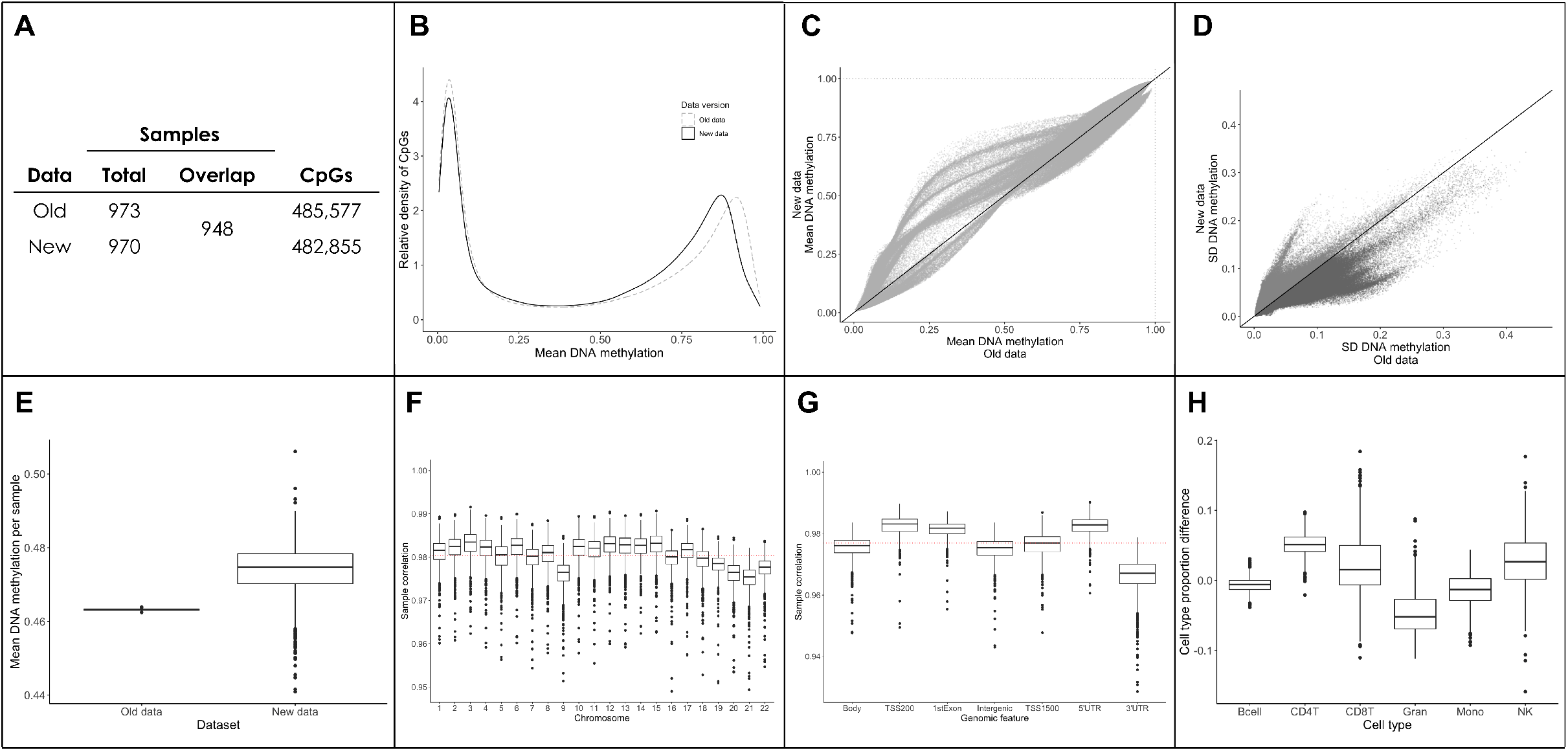
Differences between data versions of the ARIES cohort. **A)** 948 participants overlapped between versions of the data. The new dataset had slightly less probes due to filtering procedures. **B)** Both the old and the new data showed typical bimodal distributions. However, the density of genome-wide DNA methylation was shifted towards the left in the new data, suggesting that the setpoint of hypermethylated CpGs was lower in the new data. **C)** Mean values for each CpG were shifted towards more middling values in the new data. **D)** The standard deviation (SD) of each CpG was generally higher in the old data. 300,839 CpGs had higher variability in the old data (dark grey) and 182,016 CpGs had higher variability in the new data (light grey). **E)** Individual-level mean DNA methylation (across all CpGs) varied substantially between data versions. The new data were highly variable, whereas the old data showed no variability between participants. **F)** Individual-level DNAm data were generally highly correlated between data versions (r=0.98, red line), with no clear biases detected for specific chromosomes. **G)** Individual-level DNAm from specific genomic regions were generally highly correlated between data versions (r=0.98, red line). However, CpGs located in 3’UTRs showed slightly lower correlations between datasets. **H)** Estimated cell type proportions showed slight differences between the old and new datasets (differences were calculated by subtracting old data proportions from new data proportions).

For the current analyses, we further removed cross-hybridizing probes, polymorphic probes, and probes located in sex chromosomes, as well as those probes that did not overlap between both data versions. These filtering steps resulted in a list of 440,257 CpGs that were present in each data version. To remove possible outliers, we winsorized the beta values (i.e., values that represent % methylation) at each CpG site, setting the bottom 5% and top 5% of values to the 5th and 95th quantile, respectively.

### Measures of childhood adversity

We investigated seven types of childhood adversity assessed between birth and age 7: experiences of sexual/physical abuse, caregiver physical/emotional abuse, maternal psychopathology, financial stress, family instability, one-adult households, and neighborhood disadvantage. These variables were coded the same way between both the old and new datasets. For a full description of these variables, please refer to Dunn and colleagues (2019), which described their coding in depth ^5^.

## Analyses

### Epigenome-wide association study (EWAS) of childhood adversity

To determine how data versions can influence the results of traditional epigenome-wide methods, we performed EWAS for each of the childhood adversities described above using the old and new data versions. Here, we categorized children as ‘exposed’ or ‘unexposed’ to adversity on whether they experienced a given adversity between ages 0 to 7. We performed these epigenome-wide associations using the *limma* package in R ^12^. Consistent with previous work on these exposures ^5^, we included the following covariates to account for potential confounding: sex, race/ethnicity, maternal age at birth, maternal education, birth weight, number of previous pregnancies, maternal smoking during pregnancy, and cell type proportions estimated using the Houseman method ^13^. We accounted for multiple-testing using the Benjamini-Hochberg method and set the false discovery rate (FDR) at 5% ^14^.

### Structured Life Course Modeling Approach (SLCMA) of childhood adversity

The SLCMA is a two-stage method that compares different life course hypotheses that describe the relationship between time-dependent exposures and an outcome of interest ^4,15,16^. This method simultaneously compares a set of *a priori-*specified life course hypotheses encoding time-varying exposure-DNAm relationships, such as the timing of exposure (sensitive periods), or a cumulative count of exposures over time (accumulation of risk). Therefore, it provides more nuanced insights about exposure mechanisms beyond the traditional analyses of exposed versus unexposed individuals. Importantly, the SLCMA has been applied in multiple contexts to determine whether the timing of certain exposures can influence outcomes, including psychometric measures and DNAm ^3,17^. To summarize SLCMA briefly, in the first stage, variable selection (LARS-LASSO) is used to select the life course hypothesis that explains the greatest proportion of outcome variation. In the second stage, post-selection inference is performed to obtain point estimates, confidence intervals, and p-values for the hypothesis selected from the first stage, accounting for multiple testing burden associated with testing several life course hypotheses simultaneously for each locus.

To assess the impact of data version changes on SLCMA results, we tested the association between childhood adversity and epigenetic patterns, as previously reported by Dunn and colleagues (2019), in both data versions. adjusted for the same covariates as the EWAS analyses above. We tested five different life course hypotheses, including three sensitive periods hypotheses encoding exposures during the following three time periods: 1) very early childhood (0-2), 2) early childhood (3-5), 3) middle childhood (6-7); and two additive hypotheses: 4) total number exposures across childhood (accumulation), and 5) number of exposures weighted by time (recency). Post-selection inference was performed using the covariance test *(covTest)* method ^18^. We accounted for multiple-testing at the epigenome-level using the Benjamini-Hochberg method and set the FDR at 5% ^14^.

### Analytic version updates of the SLCMA of childhood adversity

To determine how updates to analytic versions influence the SLCMA results, we compared the results from the new data using the analysis described above, which we refer to as the *standard analysis*, to the latest recommendations for the SLCMA as described by Zhu and colleagues (2020), which we refer to as the *updated analysis*. This approach differed in three major ways. First, post-selection inference was performed using the selective inference method, which reduces p-value inflation compared to the covariance test in high dimensional analyses ^3,19^. Second, we adjusted for covariates using the Frisch-Waugh-Lovell (FWL) theorem (partitioned regression) ^20^. This method has been used in penalized regression analyses and can improve the statistical power to detect differences between groups ^3,21^.

Third, we updated the covariates to reflect best practices in the ALSPAC cohort, swapping parental occupation-based social class for maternal education. Maternal education is not only a better predictor of health and DNA methylation patterns, but also has better availability and comparability in other birth cohorts, allowing for more direct comparisons and integration into future meta-analyses ^22,23^.

### Sensitivity analyses of prenatal exposure to maternal smoking

Given that the associations between smoking and DNA methylation are some of the best replicated findings in the EWAS field, we performed additional sensitivity analyses to contrast this physical exposure to the psychosocial exposures described above. We assessed the impact of data versions on the association between exposure to maternal smoking during pregnancy and epigenetic patterns, as previously reported by Richmond and colleagues (2018). Following the same approach as the analyses of childhood adversity, we performed an EWAS of prenatal exposure to maternal smoking in the old and new data versions. Maternal smoking exposure was ascertained repeatedly in all three trimesters, wherein smoking at any point was considered prenatal smoking exposure ^6^. For the SLCMA analysis, we tested five separate life course hypotheses of prenatal smoking exposure: first trimester, second trimester, third trimester, accumulation across all trimesters, and recency of exposure.

## RESULTS

### Old and new versions of the ALSPAC data differed by several key descriptive features

We first assessed the CpG- and individual-level differences between the ALSPAC data normalized using the Tost pipeline (*old*) and the meffil pipeline (*new*). The genome-wide distribution of DNAm values from the old data were generally shifted towards the center in the new data (**Fig 2B and 2C**). CpG-level variability, assessed by the standard deviation of each CpG, was generally higher in the old data (**Fig 2D**). In addition, we detected higher individual-level variability (across all CpGs) in the new data than in the old data, which showed no individual-level variability due to the use of quantile normalization (**Fig 2E**). Nevertheless, individual-level data were generally highly correlated between data versions (mean r=0.981, SD=0.003), with no clear biases being detected in specific chromosomes (**Fig 2F**). However, CpGs located in 3’UTRs showed slightly lower correlations between versions (**Fig 2G**). Estimated cell-type proportions showed only slight differences between data versions but were mostly similar (**Fig 2H**).

### Epigenome-wide association study results differed between data versions

To determine how data versions may impact the results from traditional EWAS, we analyzed the association between each of the seven childhood adversity exposures and DNA methylation at age 7 in both ALSPAC DNAm data versions. Overall, we found little concordance between data versions for psychosocial exposures. In the old data, we identified one CpG at an FDR <0.05 for the abuse exposure, but no significant associations for the other adversities. By contrast, using the new data, we identified five CpGs at an FDR <0.05, but those were associated with exposure to financial stress.

Moreover, no significant CpGs overlapped between the old and new data versions (**Fig 3A**). Indeed, beyond significance thresholds, the overlap of CpGs by p-value rank was somewhat low for most adversities (10-40%) but remained higher than by random chance (**Fig 3B**).

**Figure 3.**
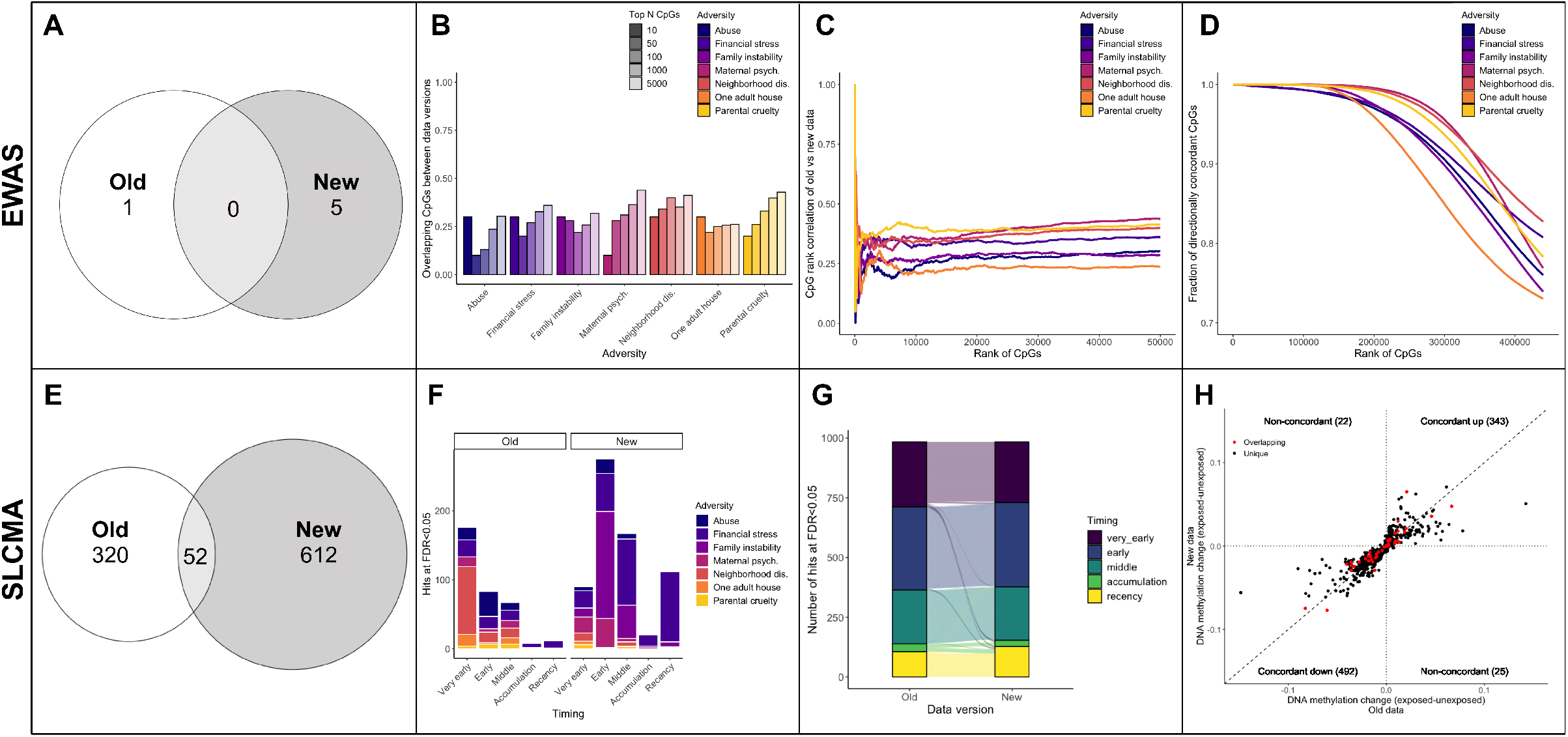
Updates to data versions change the results of epigenetic analyses, for both EWAS and SLCMA. **A)** Overlap of the hits at FDR<0.05 between the old and new data for all seven different EWAS of childhood adversity. **B)** Few CpGs overlapped between the old and new data versions at different p-value rank thresholds (top 10, 50, 100, 1000, 5000, and 50000 CpGs ranked by p-value). **C)** The Spearman’s rank correlation between CpGs (in old versus new data) that overlapped at a given rank (i.e., top N CpGs ordered by p-value) was relatively low across both data versions. **D)** The direction of DNAm differences between exposed/unexposed groups was generally consistent across overlapping CpGs at a given rank (i.e., top CpGs ranked by p-value). **E)** Overlap of the hits at FDR<0.05 between the old and new data for all seven different SLCMA of childhood adversity. **F)** Both the hypotheses selected most frequently, and the adversities identified as having the most hits varied between data versions with the SLCMA for CpGs significant at FDR<0.05. **G)** The selected hypothesis from all top hits (shown in E) were generally consistent across data versions. Each line depicted corresponds to a specific CpG and shows whether its selected hypothesis differs between analyses. **H)** The difference in DNAm values between exposed and unexposed participants across all top SLCMA hits from E was generally consistent between data versions, regardless of statistical significance. Only shown here are the CpGs associated with sensitive period hypotheses, as a the difference between exposed and unexposed individuals is not calculated for the accumulation and recency hypotheses. *Maternal psych = maternal psychopathology; Neighborhood dis = neighborhood disadvantage.

However, for each set of top CpGs (ranked by p-values), those that overlapped between data versions showed relatively good rank correlation, suggesting that some signal may be retained between data versions (**Fig 3C**). Importantly, top CpGs also showed high concordance in the direction and magnitude of differences in DNAm between exposed and unexposed groups (**Fig 3D**). As such, it appeared that the differences introduced by changing data versions caused fluctuations in the results at the level of p-value thresholds, but the results from the EWAS of childhood adversity were more similar when considering p-value ranks, as well as the direction and magnitude of associations.

### Data versions also changed the results from the SLCMA

To determine how data versions can influence more sensitive or complex methods beyond an EWAS, we assessed the impact of data versions on the SLCMA results. Here, we identified 372 CpGs in the old data and 664 CpGs in the new data at an FDR<0.05 across all seven adversities, with 52 CpGs overlapping between data versions (**Table 1**; **Fig 3E; Tables S1, S2**). The most selected hypotheses for significant CpGs were different between data versions (**Fig 3F**), as were the adversities with the most hits (**Table 1**). The old data showed more associations with *very early childhood* and neighborhood disadvantage, whereas the new data showed more associations with *early childhood* and financial stress. However, significant CpGs generally had the same hypothesis selected across data versions, with little changes in the CpGs significant in the analyses of both versions (**Fig 3G**). In addition, top hits generally showed the same direction of change and similar magnitude between data versions (**Fig 3H**). These results highlight the brittleness of p-value thresholds, which result in few overlaps between data versions, despite the general characteristics of these CpGs and their associations being similar between data versions.

**Table 1.**
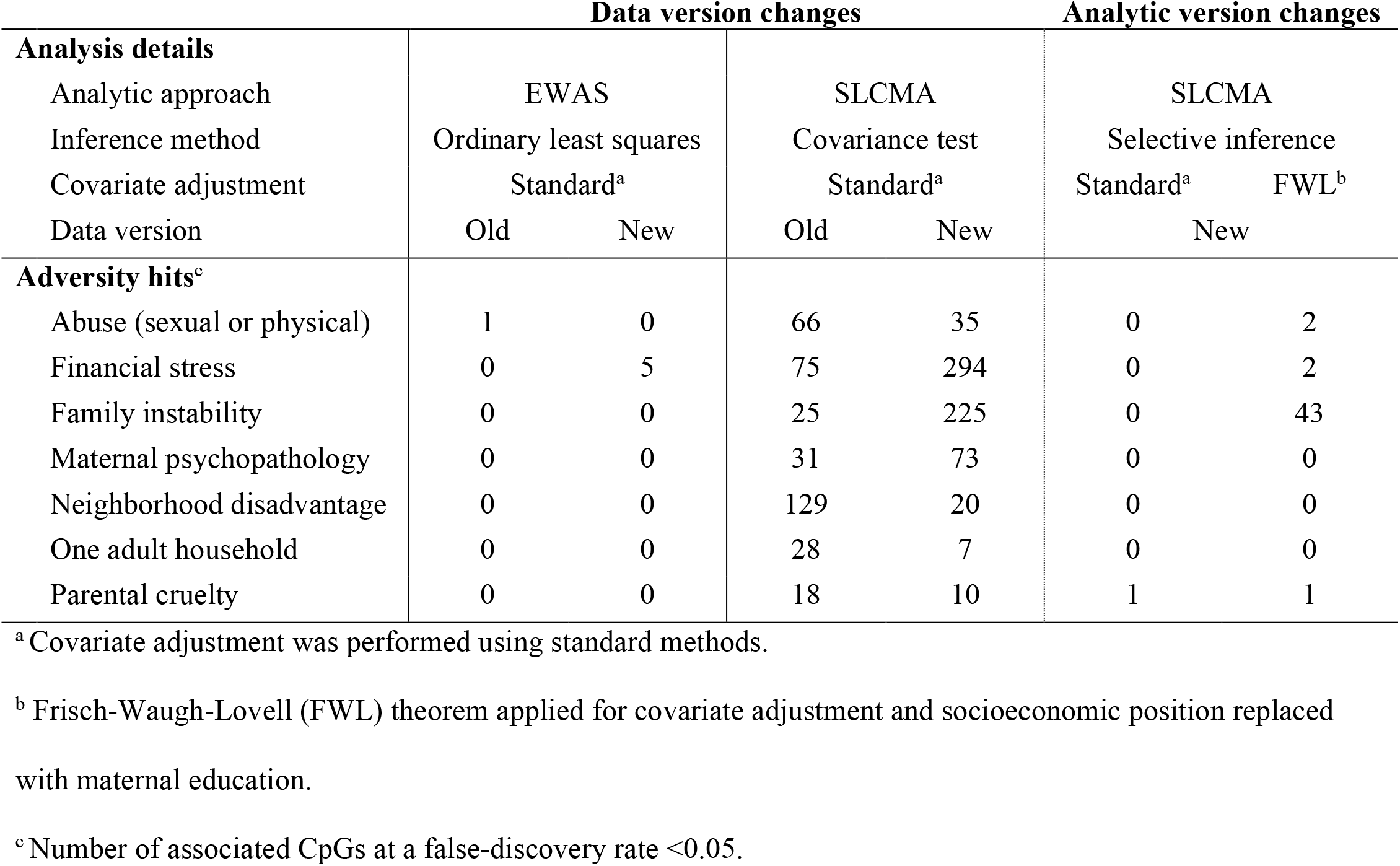
Summary of analyses and significant CpGs

### Analytic versions altered the results from the SLCMA of childhood adversity

Finally, we assessed the impact of updates to analytic versions on the results from SLCMA, as per the recommendations of Zhu and colleagues (2020) using only the new data version. We first performed the SLCMA analysis of the childhood adversities with the standard covariates and adjustment strategy but using the selective inference method in the second stage, rather than the covariance test. However, only one CpG was significant at an FDR<0.05 in this analysis. As such, we performed a comparison between the standard analytic version and the fully updated pipeline, which uses FWL correction and updated covariates. We identified 48 CpGs at an FDR<0.05 in this updated analysis, with 44 overlapping with results from the original pipeline in the new dataset (**Fig 4A; Table S3**). The majority of significant CpGs in this new analysis were association with early childhood exposure to family instability, a pattern that differed slightly from the standard version of the analysis in the new data (**Table 1**; **Fig 4B**). All significant CpGs between analytic versions showed the same hypothesis selected (**Fig 4C**). These results suggested that the reduction in power of the selective inference method can potentially be offset by the use of the FWL theorem and that updates to covariates only cause minor changes to the results. We also note that 4 CpGs overlapped between all analyses (old data with standard analysis; new data with standard analysis; new data with updated analysis), representing the associations that survived technical replication across both data and analytic versions (**Table S4**).

**Figure 4.**
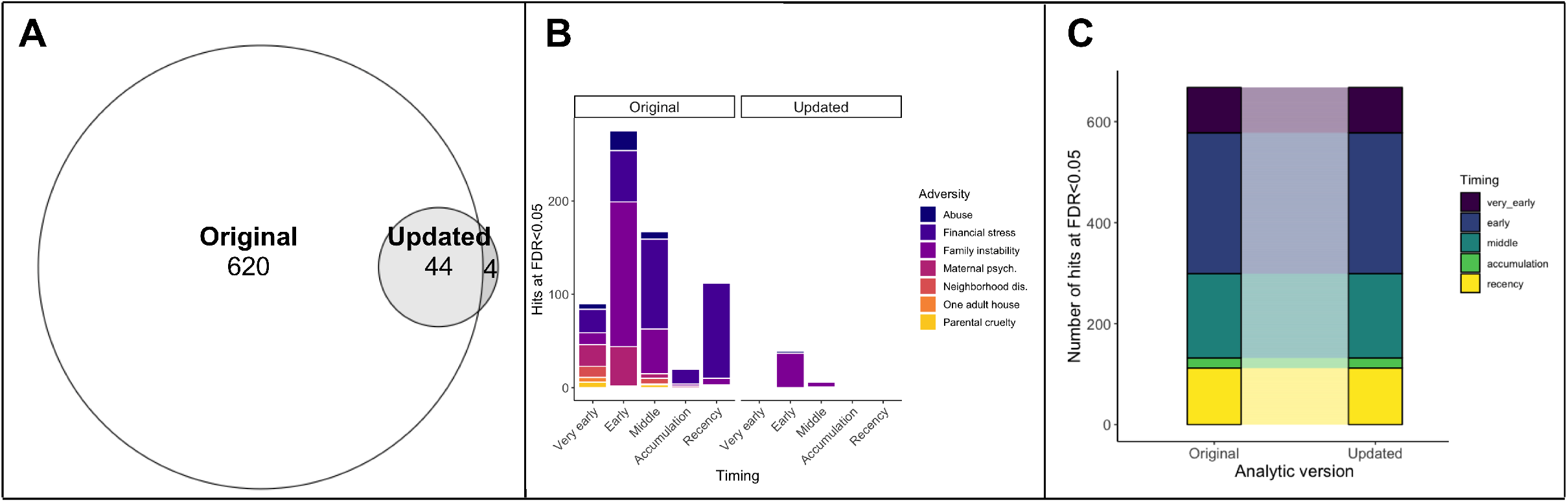
Updates to analytic versions change the results of SLCMA. **A)** Overlap of the hits at FDR<0.05 for all seven different SLCMA of adversity between the standard and updated analytic versions (analyses performed with the new data). **B)** The pattern of hypotheses selected were similar across both analytic versions, though not all adversities had statistically significant associations in the updated analytic version. **C)** The hypothesis selected across all significant CpGs from A was consistent across analytic versions. *Maternal psych = maternal psychopathology; Neighborhood dis = neighborhood disadvantage.

### Sensitivity analyses of prenatal smoke exposure showed similar results to psychosocial exposures

To determine whether the impact of data and analytic version changes were limited to psychosocial exposures, we performed secondary analyses of prenatal smoking exposure (**supplemental materials**). While the EWAS of smoking showed more overlap and consistency between data versions than psychosocial exposures (**Fig S1**), we again observed differences in terms overall concordance at the level of p-values and magnitude of change. These results suggested that p-value thresholds remain relatively arbitrary, even with “gold-standard” epigenetic associations. Our secondary analysis of prenatal smoking exposure using the SLCMA also found some overlapping CpGs at an FDR<0.05 and major changes to selected hypotheses between data versions (**Fig S2**). These results further suggest that SLCMA was more sensitive to fluctuations between data versions than EWAS, particularly during the second step of the approach when significance was assessed. We also found few overlaps between the standard and updated analytic versions of the SLCMA of prenatal smoking, suggesting that updates to covariates may have different effects on the results from SLCMA depending on analysis-specific confounding structures, since these effects were not observed with the childhood adversity analyses (**Fig S2**).

## DISCUSSION

A major challenge in conducting epigenetic analyses centers around the replicability of findings across cohorts, particularly when standard practices are constantly evolving. In this study, we quantified these differences, showing that even within the same dataset, updates to preprocessing pipelines and analytic frameworks altered the DNA methylation loci that were associated with psychosocial and physical exposures at standard p-value significance thresholds, while the magnitude of differences at these loci tended to remain the same.

The major differences between the data versions arose from two main sources: 1) individuals added or removed from the analyses due to preprocessing and withdrawal of consent for certain individuals, and 2) changes to the preprocessing pipeline for DNAm data. Although we accounted for this first factor by only analyzing overlapping samples, we found broad differences in both CpG-level and individual-level DNAm patterns that must therefore be caused by preprocessing differences. One particularly striking difference was observed at the individual level, wherein the new dataset showed increased variability across individuals due to the use of functional normalization, rather than quantile normalization in the old dataset. Such normalization techniques provide a major technical and conceptual difference in the preprocessing of DNAm data, as quantile normalization assumes that all individual samples have identical distributions of DNAm across the genome ^24^. Bulk differences between data versions were also apparent at the level of estimated cell-type proportions. Given that cell types are estimated from the DNAm data, they may reflect broader differences between data versions, which may, in turn, broadly influence the results of epigenetic analyses. Overall, no single facet of the data fully reflected the changes between datasets, suggesting that a combination of sample differences and normalization techniques likely leads to different results between versions.

As such, it is perhaps unsurprising that updates to data versions resulted in broad changes to the results of both our EWAS and SCLMA of psychosocial exposures. Although these exposures may have subtler effects on the epigenome, we found little reproducibility at the level of p-value thresholds and ranking. By contrast, the magnitude of change between exposed and unexposed individuals was highly reproducible across all CpGs in both types of analyses. For the SLCMA, we also found that hypothesis selection was stable across data versions (i.e., the first stage of SLCMA), but p-values obtained from post-selection inference were different (i.e., the second stage of SLCMA), further highlighting the fragility of inference based on p-values across our analyses. Numerous recent reports have already urged the scientific community to move away from p-values as a measure of significance and reproducibility since p-values can be less than informative and sometimes misleading ^25-28^. In particular, the American Statistical Association recently outlined six important principles to avoid the misuse of p-values in scientific analyses ^29^. They note that p-values are not a good measure of evidence on their own, nor do they measure the size or importance of an effect. Our results show these statements hold true in epigenome-wide analyses. Building from our findings and prior recommendations, we urge researchers to supplement standard analyses (e.g., reporting of p-values) with metrics that provide additional insight into the reproducibility and strength of associations, such as their magnitude and direction of effect, and allow for better understanding of both mean and variance differences within a sample^30^.

When we updated the SLCMA analytic version, we observed a not only a loss of p-value significance for several CpGs, but also several new associations. Given that we changed three main factors between analytic versions, there are at least three possible causes for these observed differences. First, selective inference is more stringent than the covariance test, which can produce inappropriately small p-values ^3^. This initial difference resulted in a total loss of FDR-significant CpGs, without any changes to the magnitude of associations, thus explaining the reduction in the number of significant CpGs. Second, the application of the FWL theorem alongside selective inference resulted in more FDR-significant CpGs. However, since the FWL theorem improves statistical power without influencing the effect estimates of associations ^3^, no new associations should arise from its application in the updated analytic version, which would explain the overlapping FDR-significant CpGs between the standard and updated analytic versions. Thus, the third difference – updates to covariates in the statistical model – is likely responsible for the emergence of four new FDR-significant CpGs in the SLCMA of psychosocial exposures. Although these differences were minor, they reflect the potential effect of moving towards more appropriate covariates in epigenome-wide analyses, such as the use of maternal education rather than occupation-based social class in the ALSPAC cohort. This result is contrasted in the secondary analyses of prenatal smoking, where changes to covariates greatly influenced the results of the analyses, highlighting that careful consideration of potential confounding is required for different types of analyses.

In contrast to the analyses of psychosocial exposures, the EWAS of prenatal smoking, a physical exposure, was relatively reproducible when using p-value thresholds. This finding was expected considering that cigarette smoke has the most reproduced findings from epigenome-wide studies ^31,32^. However, the overall ranking and overlap of CpGs beyond FDR-significance remained relatively low in the EWAS, resulting in similar levels as psychosocial exposures across the top 5,000 CpGs. These results could potentially highlight the mechanisms by which such exposures become biologically embedded. Whereas smoking exposure has not just well defined, but also targeted cellular processes (i.e., implicated pathways that clear toxins from the organism), psychosocial exposures may have more systemic influences, impacting a broader set of CpGs with smaller effects ^33,34^. In addition, it is possible that psychosocial exposures may be have greater influences in central nervous system, rather than peripheral tissues, resulting in more moderate signals from blood samples ^35^. Of note, SLCMA analyses of smoking were not well reproduced across data and analytic versions. Although these results may be due to a variety of factors, a potential explanation is that smoking may not be a time-dependent exposure. Life course modeling approaches lose power when hypotheses are highly correlated, reducing their ability to make statistical inferences ^16^. As such, these broad differences between versions may indicate that the SCLMA is not appropriate for an exposure such as prenatal smoking, which may influence epigenetic patterns equally throughout development.

The inevitable fluctuations in epigenome-wide associations highlight the importance of tracking data and analytic versions across epigenetic analyses to improve both the reproducibility and replicability of findings. As a field, we should endeavor to use the most up-to-date data versions and analytic models before performing analyses. This approach is particularly relevant for subtler exposures, such as childhood adversity, where the epigenetic signal may require more nuanced methods due to limited sample sizes. Our investigation has shown the benefit of comparing data and analytic versions in a stepwise manner (i.e. that the observed differences in results can be explained step by step). Moving beyond p-values as a single metric for significance appears to be a necessary first step towards replicability, but p-values remain an important feature of biomedical research ^28^. We propose that researchers consistently report the magnitude and direction of effects alongside p-values to provide insight into their findings. Furthermore, as CpGs tend to be highly correlated, nuanced approaches that go beyond statistical and effect size cutoffs can be used to gain broader insight into the biological mechanisms influenced by a given exposure or disease. Such methods include those assessing differentially methylated or co-methylated regions ^36,37^, or genome-wide effects, such as WGCNA and other network analyses ^38^.

## CONCLUSIONS

Changes to both data and analytic versions do impact results derived from epigenome-wide studies using both traditional and more nuanced methods. As differences not only depend on the robustness of associations, but also nuances and complexities of the analyses, our results highlight the challenges in making direct comparisons between and within datasets, stressing the importance of transparency in reporting these differences.

## ACKNOWLEDGMENTS

This work was supported by the National Institute of Mental Health of the National Institutes of Health (grant number R01MH113930 awarded to ECD). The content is solely the responsibility of the authors and does not necessarily represent the official views of the National Institutes of Health. Dr. Dunn and Dr. Lussier were also supported by a grant from One Mind. We are extremely grateful to all the families who took part in the ALSPAC study, the midwives for their help in recruiting them, and the whole ALSPAC team, which includes interviewers, computer and laboratory technicians, clerical workers, research scientists, volunteers, managers, receptionists and nurses. The UK Medical Research Council and Wellcome (Grant ref: 217065/Z/19/Z) and the University of Bristol provide core support for ALSPAC. A comprehensive list of grants funding is available on the ALSPAC website (http://www.bristol.ac.uk/alspac/external/documents/grant-acknowledgements.pdf); This research was specifically funded by grants from the BBSRC (BBI025751/1; BB/I025263/1), IEU (MC_UU_12013/1; MC_UU_12013/2; MC_UU_12013/8), National Institute of Child and Human Development (R01HD068437), NIH (5RO1AI121226-02), and CONTAMED EU (212502). This publication is the work of the authors, whom will serve as guarantors for the contents of this paper. Dr. Walton is funded by CLOSER, whose mission is to maximise the use, value and impact of longitudinal studies. CLOSER was funded by the Economic and Social Research Council (ESRC) and the Medical Research Council (MRC) between 2012 and 2017. Its initial five-year grant has since been extended to March 2021 by the ESRC (grant reference: ES/K000357/1). The funders took no role in the design, execution, analysis or interpretation of the data or in the writing up of the findings. www.closer.ac.uk. Dr. Walton is also supported by the European Union’s Horizon 2020 research and innovation programme (grant n° 848158).

