## Supplemental tables and figures for "Updates to data versions and analytic methods influence the reproducibility of results from epigenome-wide association studies"

**Table S1.** Summary of analyses of prenatal smoking and significant CpGs at FDR<0.05

| | **Data version changes** | | | | **Analytic version changes** | |
|---|---|---|---|---|---|---|
| **Analysis details** | | | | | | |
| Analytic approach | EWAS | | SLCMA | | SLCMA | |
| Inference method | Ordinary least squares | | Covariance test | | Selective inference | |
| Covariate adjustment | Standard[#] | | Standard[#] | | Standard[#] | FWL* |
| Data version | Old | New | Old | New | New | |
| Prenatal smoking hits[&] | 27 | 23 | 25 | 4576 | 0 | 13 |

[#]Covariate adjustment was performed using standard methods.
*Frisch-Waugh-Lovell (FWL) theorem applied for covariate adjustment and socioeconomic position replaced with maternal education.
[&] Number of associated CpGs at a false-discovery rate <0.05.

### SUPPLEMENTAL FIGURES

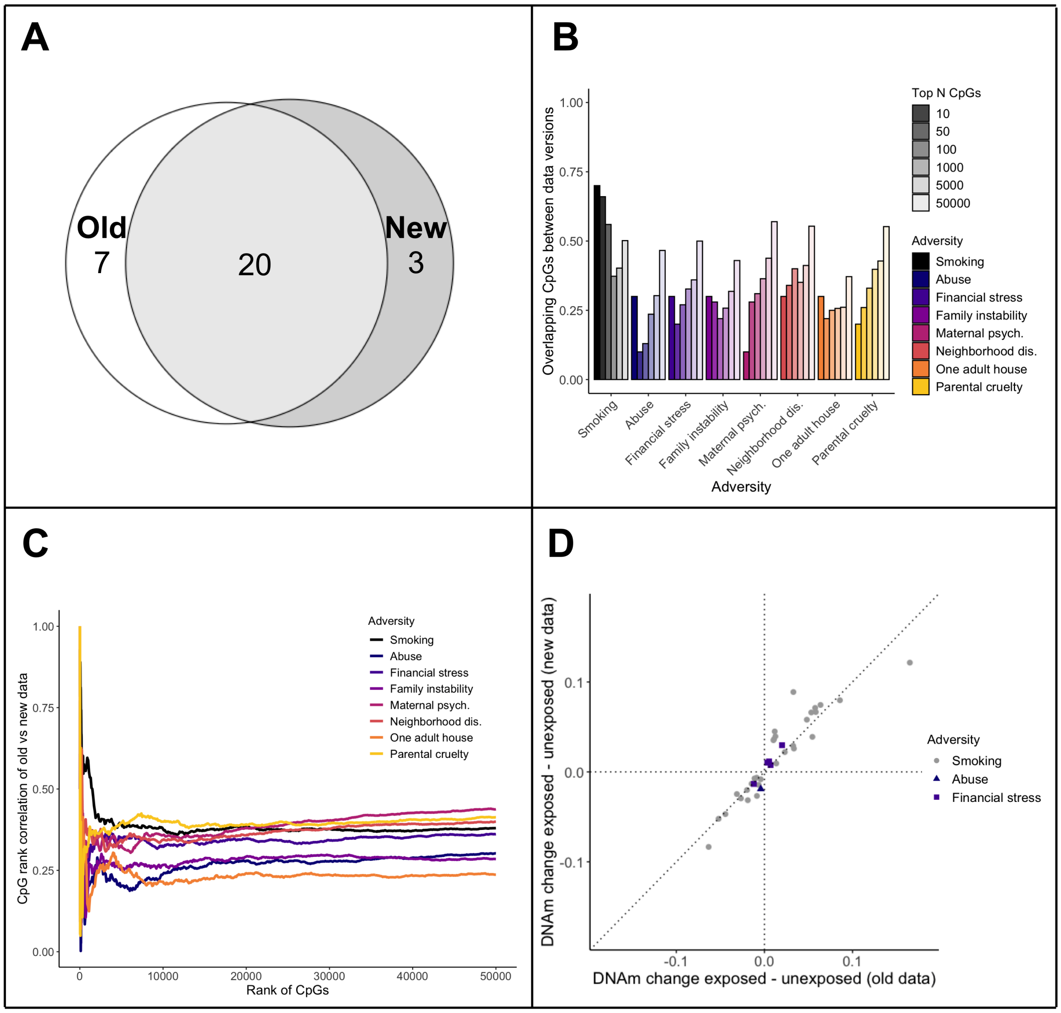


**Figure S1. Results from the EWAS of prenatal smoking and compared to the EWAS of postnatal adversity.**

**A)** Overlap of the hits at FDR<0.05 for the EWAS of prenatal smoking exposure between the old and new data.

**B)** Few CpGs overlapped between data versions at different rank thresholds for the adversities (top 10, 50, 100, 1000, and 5000 CpGs ordered by p-value). However, prenatal smoking showed higher overlaps between top ranked CpGs.

**C)** The Spearman's rank correlation between CpGs (in old versus new data) that overlapped at a given rank (i.e., top N CpGs ordered by p-value) was relatively low across both data versions.

**D**) The direction of change between exposed and unexposed groups was consistent for all significant CpGs in both prenatal smoking and postnatal adversity (abuse, financial stress).

*Maternal psych. = maternal psychopathology; Neighborhood dis. = neighborhood disadvantage.

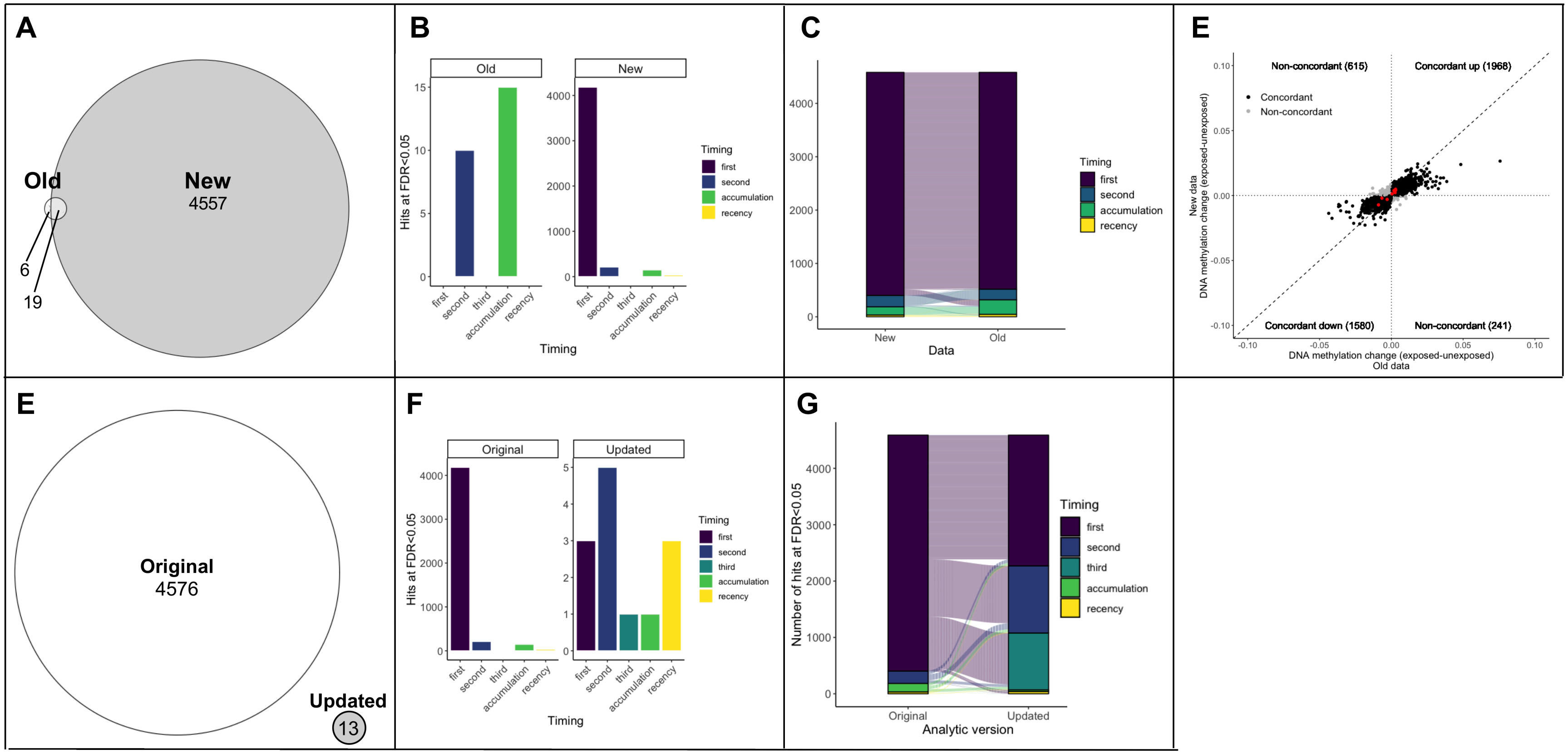


**Figure S2. Results from the SLCMA of prenatal smoking.**

**A)** Overlap of the hits at FDR<0.05 for the SLCMA of prenatal smoking between the old and new data.

**B)** The hypotheses selected most frequently across SLCMA hits were different between data versions (note that the scales are different between the panels of B).

**C)** The selected hypothesis of all top hits from E were generally consistent across analyses. Here, each line is a given CpG and shows how its selected hypothesis changes between analyses.

**D)** The change in DNAm between exposed and unexposed individuals across all top SLCMA hits from E was generally consistent between data versions, regardless of significance (red = overlapping CpGs from A).

**E)** Overlap of the hits at FDR<0.05 for the SLCMA of prenatal smoking between the standard and updated analytic versions (new data).

**F**) Different patterns of hypothesis selected were present across both analytic versions (note that the scales are different between the panels of F).

**G)** The hypothesis selected across all significant CpGs from E was generally different across analytic versions.
